# Workflows for detecting fungicide resistance in net form and spot form net blotch pathogens

**DOI:** 10.1101/2023.04.27.538624

**Authors:** N. L. Knight, K. C. Adhikari, K. Dodhia, W. J. Mair, F. J. Lopez-Ruiz

## Abstract

Fungicide resistance in *Pyrenophora teres* f. *maculata* and *P. teres* f. *teres* has become an important disease management issue. Control of the associated barley foliar diseases, spot form and net form net blotch, respectively, relies on three major groups of fungicides, demethylation inhibitors (DMI), succinate dehydrogenase inhibitors (SDHI) and quinone outside inhibitors (QoI). However, resistance has been reported for the DMI and SDHI fungicides in Australia. To enhance detection of different resistance levels, phenotyping and genotyping workflows were designed. The phenotyping workflow generated cultures directly from lesions and compared growth on discriminatory doses of tebuconazole (DMI) and fluxapyroxad (SDHI). Genotyping real-time PCR assays were based on alleles associated with sensitivity or resistance to the DMI and SDHI fungicides. These workflows were applied to a net blotch collection from 2019 consisting predominantly of *P. teres* f. *teres* from South Australia and *P. teres* f. *maculata* from Western Australia. For South Australia the *Cyp51A* L489-3 and *SdhC*-R134 alleles, associated with resistance to tebuconazole and fluxapyroxad, respectively, were the most prevalent. These alleles were frequently found in single isolates with dual resistance. This study also reports the first detection of a 134 base pair insertion located at position −66 (PtTi-6) in the *Cyp51A* promoter of *P. teres* f. *maculata* from South Australia. For Western Australia, the PtTi-1 insertion was the most common allele associated with resistance to tebuconazole. These workflows will be valuable for screening *P. teres* populations for fungicide resistance, and informing appropriate management strategies.

## Introduction

Fungicides are a necessary disease management tool in many cropping systems, however the efficiency of disease control is jeopardised by the emergence of decreased sensitivity to fungicides in fungal pathogen populations. Decreased sensitivity to fungicides may be characterised by *in vitro* screening or by observing a loss of efficacy of fungicide treatments in the field, with responses across increasing fungicide concentrations ranging from sensitivity to reduced sensitivity to resistance (Ireland et al. 2021). Monitoring for decreased fungicide sensitivity uses similar processes, particularly phenotypic assessments of fungal isolates on fungicide amended media and targeted gene assessment. Once genotypes associated with reduced sensitivity or resistance are known, assays can be designed for DNA-based detection directly from mycelia or diseased tissues (Hellin et al. 2020). This information is critical for informing disease management strategies.

In Australia, fungicide resistance in the barley (*Hordeum vulgare*) pathogen *Pyrenophora teres* has become an important management issue (Ellwood et al. 2019; Mair et al. 2016; Mair et al. 2020; Mair et al. 2023). *Pyrenophora teres* is recognised as two forms, *P. teres* f. *teres* and *P. teres* f. *maculata*, causing net form net blotch (NFNB) and spot form net blotch (SFNB), respectively (Smedegård-Petersen 1971). Hybridisation between the two forms is also possible (Campbell et al. 2002; Campbell and Crous 2003). Net blotch has been reported to cause yield losses of up to 36.5% for NFNB and 44% for SFNB, with the potential to cause even greater losses under favourable environmental conditions (Jayasena et al. 2007; Khan 1987; Murray and Brennan 2009; Steffenson et al. 1991). These stubble-borne pathogens are managed using crop rotations, fungicide applications and host genetic resistance (McLean et al. 2009; Wallwork 2000). Resistant cultivars are the preferred option due to the costs associated with fungicide application, however the genetic variation present in the *P. teres* f. *teres* and *P. teres* f. *maculata* populations may overcome available resistance sources (Akhavan et al. 2017; Boungab et al. 2012; Fowler et al. 2017; Gupta et al. 2012; Platz et al. 2000; Wu et al. 2003).

Fungicide groups available for the control of the net blotch diseases include demethylation inhibitors (DMI; FRAC group 3), succinate dehydrogenase inhibitors (SDHI; FRAC group 7) and quinone outside inhibitors (QoI; FRAC group 11) (Ireland et al. 2021). These include a range of specific chemicals, however registrations for net blotch control can vary by country and region. While these fungicide groups can provide disease control for sensitive fungal populations, reduced sensitive or resistant types pose a risk (Olvång 1988). Reduced sensitivity or resistance in *P. teres* has been described for DMI (Campbell and Crous 2002; Ellwood et al. 2019; Mair et al. 2016; Mair et al. 2020; Sheridan et al. 1985; Sheridan and Grbavac 1985), SDHI (Mair et al. 2023; Rehfus et al. 2016) and QoI fungicides (Marzani et al. 2013; Semar et al. 2007; Sierotzki et al. 2007).

Genotypes of *P. teres* associated with sensitivity, reduced sensitivity and resistance have been reported for DMI, SDHI and QoI fungicides. These include single nucleotide polymorphisms (SNPs) in the genes for cytochrome P450 sterol 14α-demethylase (CYP51A) (Mair et al. 2016), the succinate dehydrogenase (SDH) B, C and D sub-units (Mair et al. 2023; Rehfus et al. 2016) and cytochrome b (Sierotzki et al. 2007), along with a range of 134 bp insertions in the *Cyp51A* promoter region (Mair et al. 2020). Detection of these genotypes may be achieved using methods such as pyrosequencing (Gobeil-Richard et al. 2016; Santos et al. 2020) and real-time or digital PCR (Capote et al. 2012; Hellin et al. 2020; Zulak et al. 2018). Each method provides relevant information, however the relative commonality of PCR infrastructure across laboratories and the low limit of detection makes real-time or digital PCR-based detection an effective option.

PCR-based detection and quantification of SNP genotypes requires discrimination at a single base position (Bottema and Sommer 1993; Newton et al. 1989). Allele-specific PCR design methods must consider the different effects of mismatched bases on amplification efficiencies (Huang et al. 1992; Stadhouders et al. 2010) and may include specific positioning of target sequences at the 3′ end in one primer (Newton et al. 1989) or the inclusion of locked nucleic acids on or adjacent to the SNP within a hydrolysis probe or primer (Hellin et al. 2020; You et al. 2006) to yield adequate discrimination. Billard et al. (2012) described a strategy encompassing both primer and probe alignment with a single SNP, which harnesses the discriminatory functions of both primer and probe mismatches for non-target DNA, while allowing efficient amplification of target sequences. Inclusion of deliberate artificial mismatches in the primer sequence may further improve allelic discrimination (Billard et al. 2012; De Miccolis Angelini et al. 2014; Glaab and Skopek 1999; Newton et al. 1989).

Informed disease management decisions in the field are best supported by a combination of phenotypic and genotypic detection strategies for fungicide resistance. The identification of reduced sensitivity and resistance to DMI and SDHI fungicides in *P. teres* f. *maculata* and *P. teres* f. *teres*, and characterisation of associated alleles, demonstrates the need for comprehensive detection methods. This study aimed to outline a workflow for efficiently phenotyping *P. teres* isolates from net blotch affected barley leaves on DMI and SDHI amended media, followed by genotyping of isolates. Real-time PCR genotyping assays were designed to include *P. teres* form-specific identification and allele-specific genotyping of sequences associated with sensitivity, reduced sensitivity and resistance phenotypes. The phenotyping and genotyping workflows were used to assess a collection of net blotch affected leaf samples predominantly from South Australia and Western Australia.

## Materials and methods

### Isolation and fungicide resistance test

Net blotch affected barley leaves and *P. teres* isolates were provided by growers, agronomists and researchers in New South Wales, South Australia, Victoria, and Western Australia in 2019. Leaf sampling was also performed by the authors in South Australia and Western Australia.

Leaf samples were surface sterilised by washing for 30 s in 70% (v/v) ethanol, 60 s in 0.125% (w/v) NaOCl and 2× 60 s in autoclaved filtered water. After air drying, lesioned tissue sections (including lesion edges) of approximately 0.5 cm^2^ were removed and inserted into potato dextrose agar (PDA) with antibiotics (39 g/L PDA + 0.1 g/L ampicillin, 0.05 g/L neomycin and 0.03 g/L streptomycin). Two millilitres of agar media were dispensed in each well of a 12 well plate.

After seven days of incubation at room temperature under natural light conditions, fungal growth was assessed. Mycelium of cultures determined to be *P. teres*, based on colony colour and morphology, was scraped from the media and suspended in 2 mL tubes containing two 3 mm diameter steel ball bearings and 500 µL of autoclaved filtered water. Hyphal fragments were generated by milling at 30 Hz for 2× 10 s (Mixer Mill MM 400, Retsch GmbH, Haan, Germany). A 10 µL aliquot of the hyphal suspension was placed onto 1 mL of control or fungicide amended media in a 24 well plate and incubated in the dark at room temperature. Fungicide amended media included tebuconazole (DMI; 0 µg/mL, 15 µg/mL and 50 µg/mL) in PDA or fluxapyroxad (SDHI; 0 µg/mL, 2 µg/mL, 5 µg/mL and 10 µg/mL) in yeast bacto acetate agar (YBA; Mair et al. 2020). Growth was assessed at seven days after plating on each discriminatory dose. Sub-cultures were taken from mycelium growing on the greatest fungicide concentration and placed onto V8-potato dextrose agar (V8-PDA; Mair et al. 2016).

Single-spore cultures were generated by placing 0.5 cm^2^ plugs of seven-day-old V8-PDA cultures upside down on water agar (15 g/L agar) and incubating for up to six days at 22 °C under a 12-hour photoperiod of white light. Single spores were picked with an acupuncture needle and transferred to V8-PDA. Growth of each pure culture was assessed on fungicide amended media as described above.

### DNA extraction of isolate collection

For initial PCR genotyping optimisation, DNA of standard isolates (Supplementary Table 1) and barley was required. Briefly, mycelia from standard isolates (cultures grown for 14 days on V8-PDA) or barley leaves (seven-day-old barley seedlings [cv. Baudin] grown at room temperature with a 12-h photoperiod) were ground with two 3 mm diameter steel balls at 30 Hz for 60 s (Mixer Mill MM 400). DNA was extracted using a BioSprint 15 DNA Plant Kit (Qiagen, Hilden, Germany) according to the manufacturer’s instructions and stored at −20 °C. DNA concentrations were measured with a Qubit Flex Fluorometer (Thermo Fisher Scientific, Waltham, MA, USA) using a Qubit dsDNA BR assay kit (Thermo Fisher Scientific).

For genotyping, DNA of the entire collection was extracted from mycelia of cultures grown for 14 days on V8-PDA following the method described by Dodhia et al. (2021). Briefly, harvested mycelia was placed in a 2 mL tube containing two 3 mm steel ball bearings and 400 µL of lysis buffer, ground at 30 Hz for 60 s (Mixer Mill MM 400) and centrifuged at 20627 × *g* for 5 min. Ten microliters of supernatant was then diluted into 990 µL of 1× TE buffer. DNA samples were stored at −20 °C prior to assessment in real-time PCR.

### *Pyrenophora teres* form-specific PCR design

The design of a duplex PCR assay for simultaneous specific identification of *P. teres* f. *maculata* and *P. teres* f. *teres* was based on form-specific DNA regions described by Mair et al. (2023). Briefly, a hydrolysis probe sequence was positioned between the forward and reverse primer positions and assessed for oligonucleotide characteristics in Primer3 v. 0.4.0 (Koressaar and Remm 2007; Untergasser et al. 2012) and sequence similarity by a standard nucleotide Basic Local Alignment Search Tool (BLAST) search of the GenBank Standard databases (nr etc.) (Altschul et al. 1990). PCR optimisation and specificity assessment was then performed using real-time PCR and DNA of standard isolates. The DNA templates included 2.5 ng of single target or non-target DNA and mixed samples containing 2.5 ng of DNA template of isolates SG1 and 9254. Reactions for uniplex and duplex assays were performed in 20 μL volumes containing 10 μL Immomix (Bioline) and 5 μL of DNA template. Each reaction contained 0.15 μM of each probe (Integrated DNA Technologies) and 0.25 μM of each forward and reverse primer (Integrated DNA Technologies; Table 1). Thermal cycling consisted of an initial denaturation for 10 min at 95 °C followed by 40 cycles of 95 °C for 15 s and 60 °C for 60 s. Real-time PCR was performed in a CFX96 or CFX384 real-time system (Bio-Rad Laboratories, Hercules, CA, USA). Fluorescence data for each fluorophore was collected during the 60 °C step of each cycle and visualised in the CFX Maestro Software v. 1.1 (Bio-Rad Laboratories).

**Table 1.**
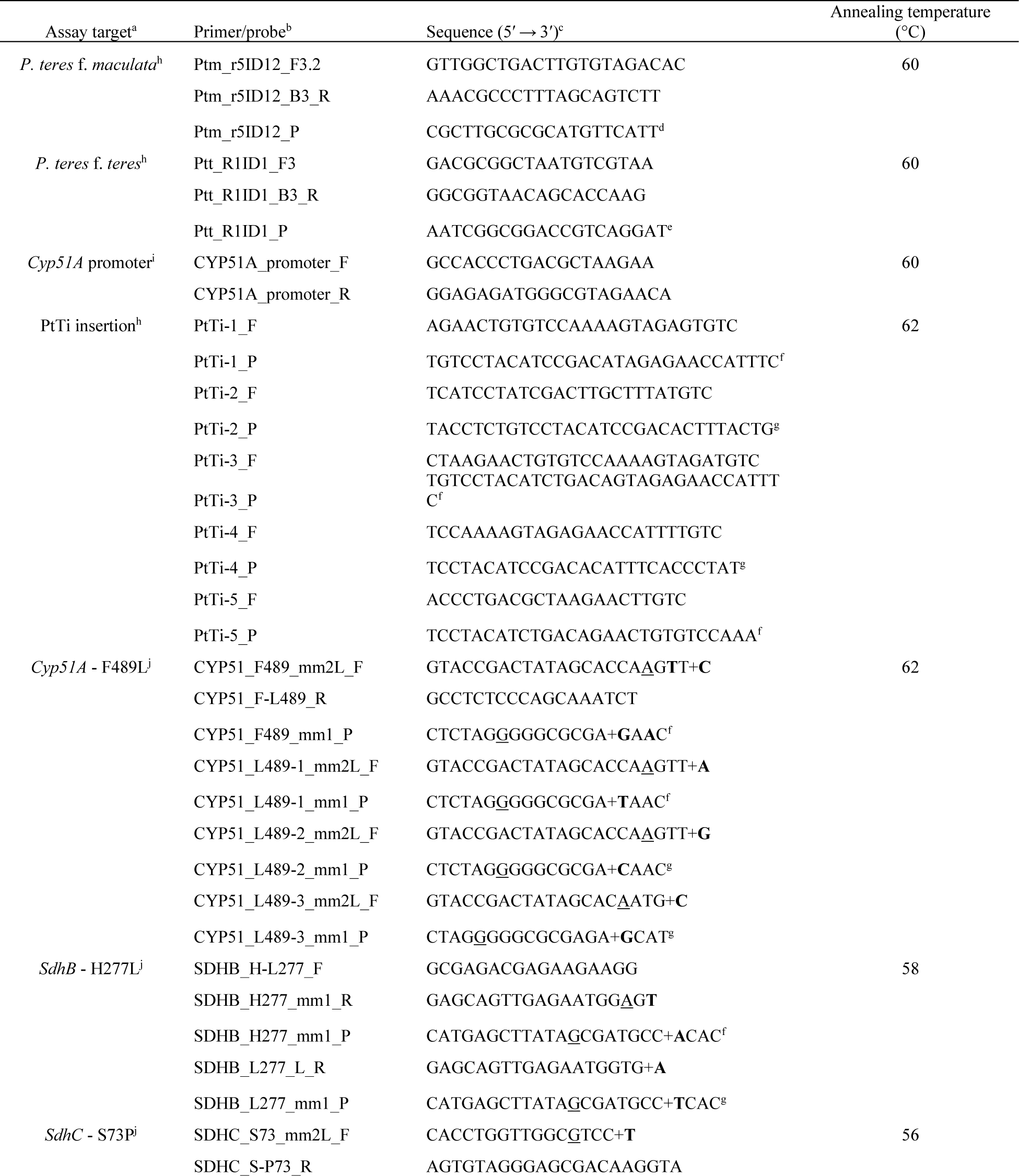
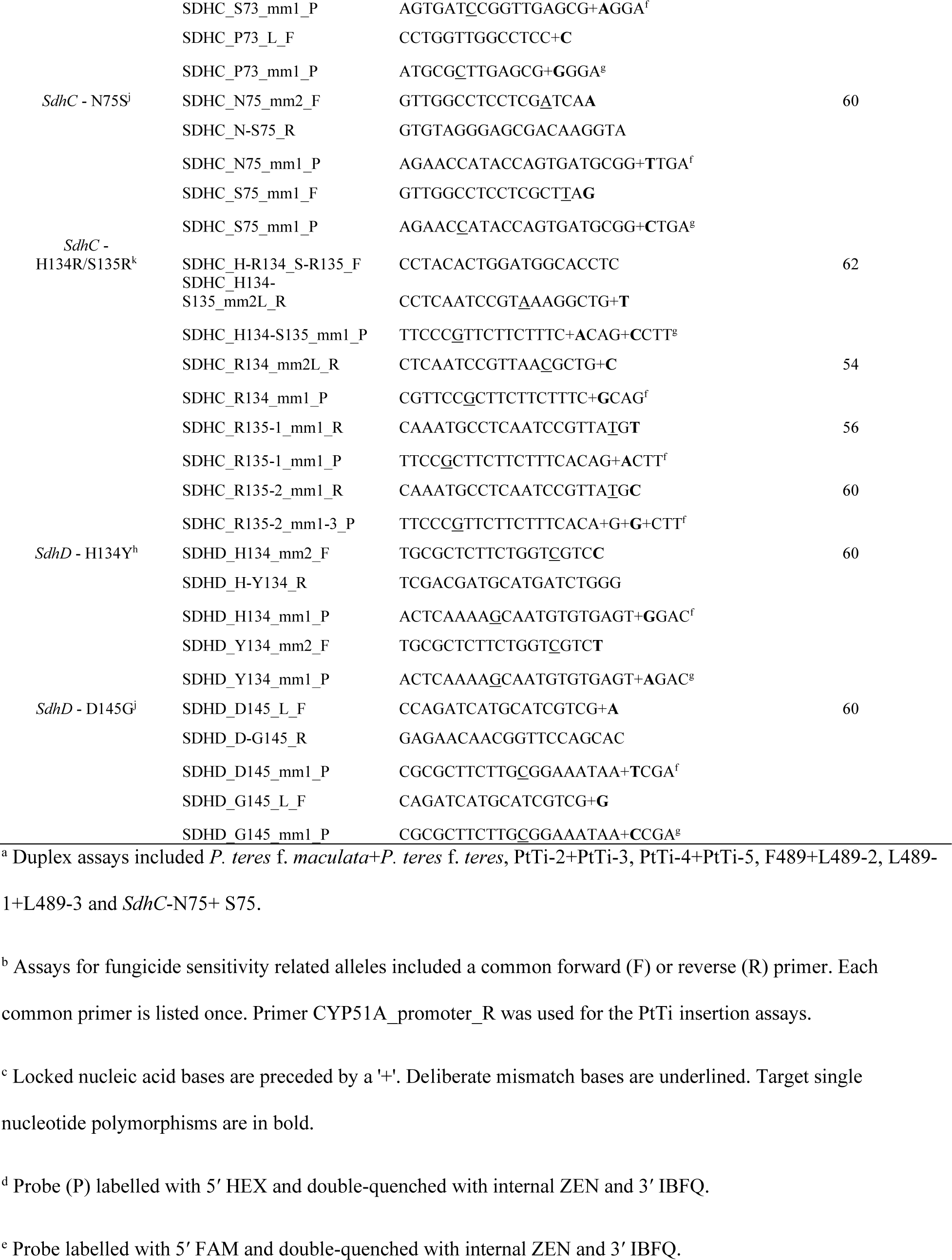
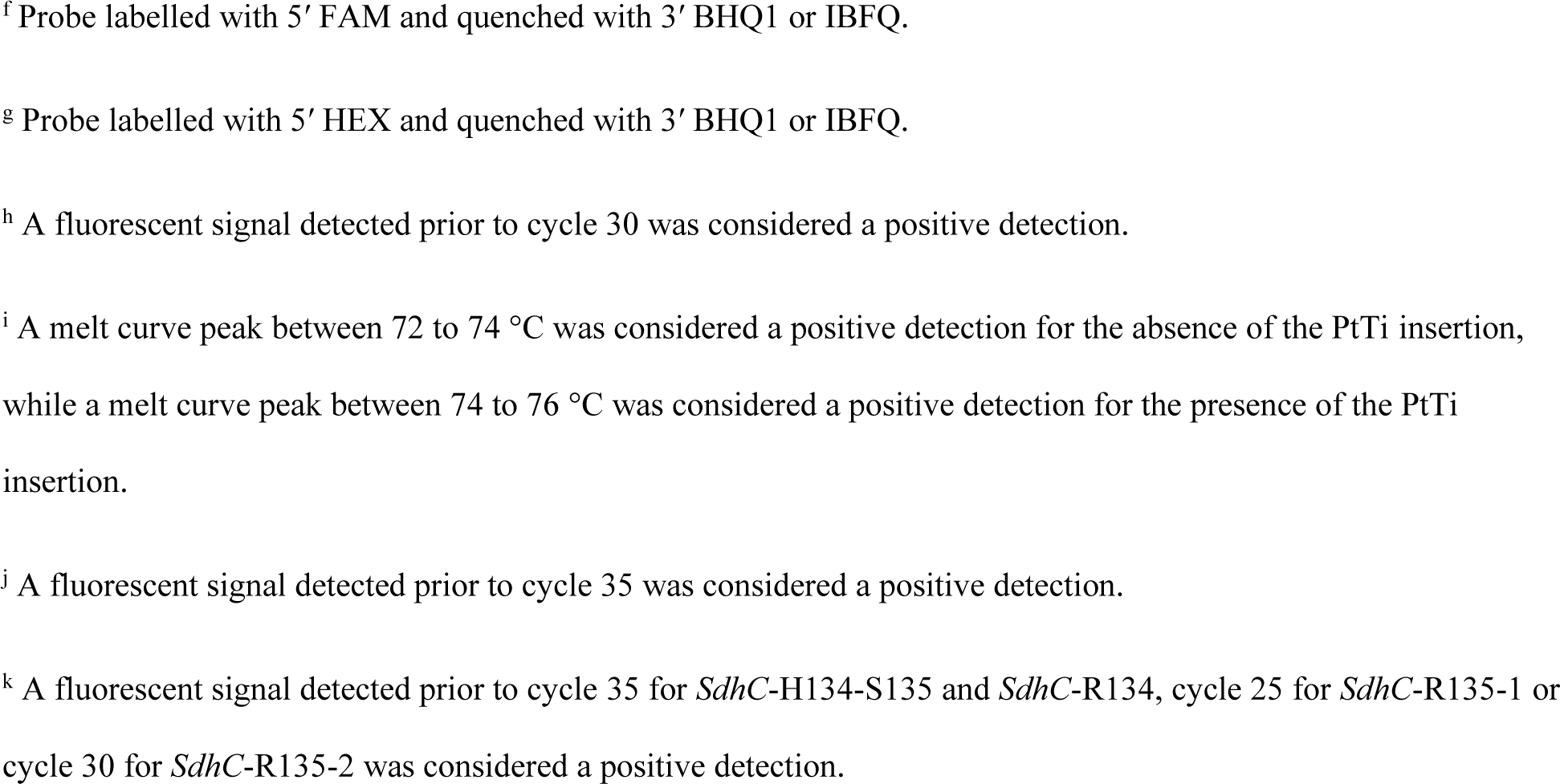
Primer/probe sets for detection of *P. teres* f. *maculata*, *P. teres* f. *teres* and fungicide sensitivity related genotypes.

### Allele-specific PCR design

Hydrolysis probe real-time PCR assays were designed for alleles of the *Cyp51A* promoter and coding regions, and the *SdhB*, *SdhC* and *SdhD* subunit genes associated with sensitive, reduced sensitive and resistant phenotypes (Table 1). Primer and probe characteristics were assessed using Primer3 v. 0.4.0 (Koressaar and Remm 2007; Untergasser et al. 2012). All real-time PCR reactions were performed in 20 μL volumes containing 10 μL Immomix (Bioline) and 5 μL of DNA template, with thermal cycling conducted in a CFX96 or CFX384 real-time system (Bio-Rad). Primer and probe concentrations are provided in the descriptions below. Fluorescence data was visualised in the CFX Maestro Software v. 1.1 (Bio-Rad Laboratories).

Optimisation of each assay initially included uniplex gradient real-time PCR of target and non-target DNA templates, with optimal discrimination determined by comparing the quantification cycles (Cq) of target and non-target DNA. This was followed by uniplex and duplex real-time PCR with target, non-target, combined target/non-target, negative control (barley) and no template control (water) templates. DNA quantities for each target/non-target were constant within an assay (either 2.5 or 0.25 ng) to allow comparison of fluorescence detection.

Assays for the *Cyp51A* promoter PtTi insertion alleles were designed to detect either the presence or absence of the PtTi insertion or the specific PtTi-1 to 5 alleles. The assay detecting the presence or absence of the *Cyp51A* promoter insertion was designed by placing primers on conserved regions either side of the insertion positions (Supplementary Figure 1a) and using melt curves to discriminate between short (86 bp – no insertion) and long (220 bp – with insertion) PCR products. Each reaction contained 0.5 μM of each forward and reverse primer (Integrated DNA Technologies) and 1× SYBR Green (Invitrogen). Thermal cycling consisted of an initial denaturation for 10 min at 95 °C followed by 40 cycles of 95 °C for 15 s, 60 °C for 30 s and 72 °C for 30 s. PCR product dissociation was measured from 65 °C to 80 °C at 0.1 °C increments for 5 s. Fluorescence data was collected during the 72 °C step of each cycle and across the melting gradient to generate melt curves.

Assays for specifically detecting PtTi-1 to 5 insertions included a forward primer and a probe, respectively, across the 5′ and 3′ boundaries of the PtTi sequence within the *Cyp51A* promoter region (Supplementary Figure 1b). A reverse primer designed to a conserved region was used for each assay (Table 1). Each uniplex reaction contained 0.15 μM of probe (Integrated DNA Technologies; Table x) and 0.25 μM of each forward and reverse primer (Integrated DNA Technologies). Duplex assays contained the same concentrations for each probe and forward primer, while the reverse primer was 0.5 μM. Thermal cycling consisted of an initial denaturation for 10 min at 95 °C followed by 40 cycles of 95 °C for 15 s and 62 °C for 60 s. Fluorescence data was collected during the 62 °C step of each cycle.

PCR-based discrimination of SNP alleles in the *Cyp51A*, *SdhB*, *SdhC* and *SdhD* genes was investigated using modifications to the process described by Billard et al. (2012). Briefly, forward primers were designed to align to the SNP at the 3′ terminus (Petruska et al. 1988). A probe was positioned to partially overlap with the forward primer binding position in the opposite strand, which included the target SNP. A reverse primer designed to a conserved region was used for each allele assay (Table 1; Figure 1). The SNP binding positions in the forward primers and probes were assessed with conventional bases or locked nucleic acids, which may increase allele-specific PCR specificity (Latorra et al. 2003; You et al. 2006) through altered binding dynamics. Deliberate artificial mismatches were also included into modified forward primers and probes to improve hybridisation discrimination. The selection of the base identity for artificial mismatches was informed by mismatch effects reported by Stadhouders et al. (2010). Optimal discrimination was determined by comparing the quantification cycles (Cq) of target and non-target DNA of the same concentration. Primer and probe concentrations for uniplex and duplex reactions, and thermal cycling, were as described above with annealing temperatures assay dependent (Table 1).

**Figure 1.**
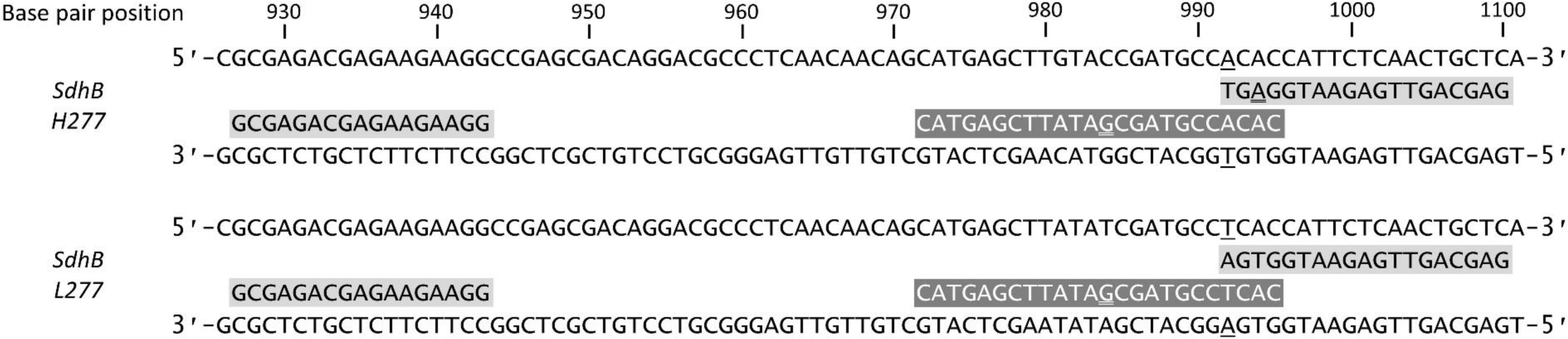
Example of the primer and probe alignment method for single nucleotide polymorphism (SNP) discrimination in real time PCR. This demonstrates the alignment of primers (black text on grey) and probes (white text on grey) against the succinate dehydrogenase subunit b (*SdhB*) gene sequences (black text on white) of *Pyrenophora teres* containing the *SdhB*-H277 or *SdhB*-L277 allele. Underlined bases in the gene sequence were the single nucleotide polymorphisms targeted for specific detection. Underlined bases in the primer or probe sequences were deliberate mismatches. GenBank accessions for each sequence were KU534598 (*SdhB*-H277) and ON534228 (*SdhB*-L277). GenBank accession number KU534598 was used for base pair reference positions.

### Real-time PCR genotyping

DNA of each isolate was genotyped in real-time PCR using the form-specific and allele-specific assays described above. DNA templates were run in duplicate and each PCR plate included positive and negative control templates for the respective assays. A mean Cq value was used to determine each genotype. Isolates exhibiting a reduced sensitive or resistant phenotype to fungicides but no corresponding genotype were re-tested.

### Sequencing

Isolates with a resistance phenotype but no detectable resistance genotype were sequenced for the corresponding genes. These included the *Cyp51A* promoter of eight isolates positive for the PtTi insertion but no detectable position and the *SdhB*, *C* and *D* sub-units for one isolate exhibiting a FLUX-10 resistance phenotype. Sequencing was performed as previously described (Mair et al. 2020; Mair et al. 2023).

## Results

### Fungicide resistance phenotypes

The *P. teres* collection consisted of 258 isolates (Supplementary Table 1). This included 17 standard isolates, and 246 isolates collected in 2019. The 2019 collection originated from New South Wales (*n* = 5, *P. teres* f. *maculata*), South Australia (*n* = 160, with 30 *P. teres* f. *maculata* and 130 *P. teres* f. *teres*), Victoria (*n* = 6, *P. teres* f. *teres*) and Western Australia (*n* = 75, with 73 *P. teres* f. *maculata* and 2 *P. teres* f. *teres*).

The discriminatory doses of tebuconazole and fluxapyroxad identified phenotypes linked to reduced sensitivity (15 µg/mL tebuconazole and 2 or 5 µg/mL of fluxapyroxad) and resistance (50 µg/mL tebuconazole and 10 µg/mL of fluxapyroxad) (Table 2). All NSW isolates were sensitive to both tebuconazole and fluxapyroxad, while two *P. teres* f. *teres* from Victoria were resistant to tebuconazole. For the South Australia collection, 20% of the *P. teres* f. *maculata* were reduced sensitive, while 21% of the *P. teres* f. *teres* were reduced sensitive and 49% were resistant to tebuconazole. In comparison, all the South Australian *P. teres* f. *maculata* were sensitive to fluxapyroxad, while 8% of the *P. teres* f. *teres* were reduced sensitive and 48% were resistant. Dual resistance phenotypes were also detected in *P. teres* f. *teres* from South Australia, with 3% reduced sensitive to both tebuconazole and fluxapyroxad, 12% reduced sensitive to tebuconazole and resistant to fluxapyroxad, 2% resistant to tebuconazole and reduced sensitive to fluxapyroxad, and 32% resistant to both tebuconazole and fluxapyroxad. For the Western Australia collection, 52% of the *P. teres* f. *maculata* were reduced sensitive and 26% were resistant to tebuconazole, while the *P. teres* f. *teres* consisted of one reduced sensitive and one resistant phenotype. All the Western Australia isolates were sensitive to fluxapyroxad.

**Table 2.**
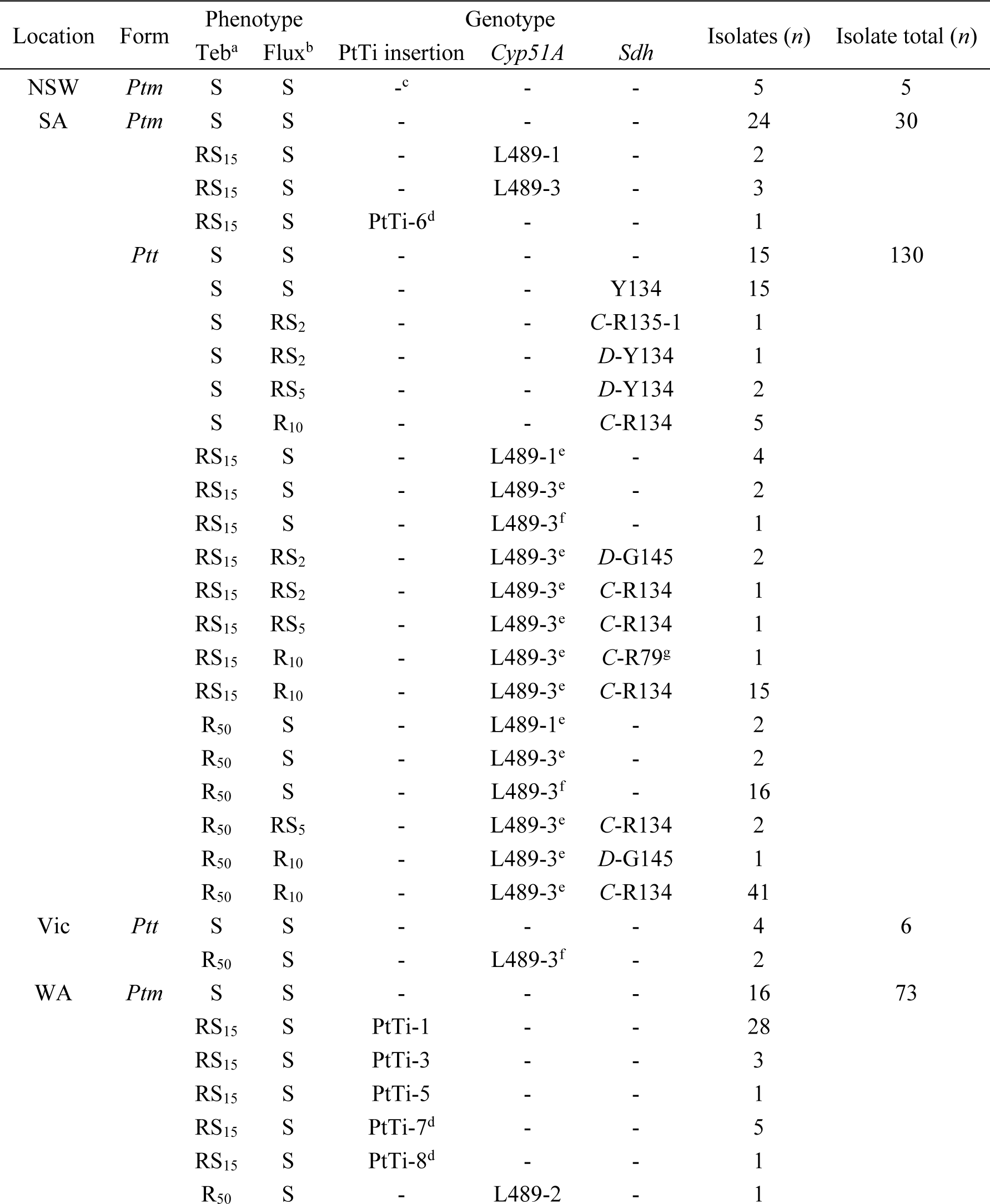
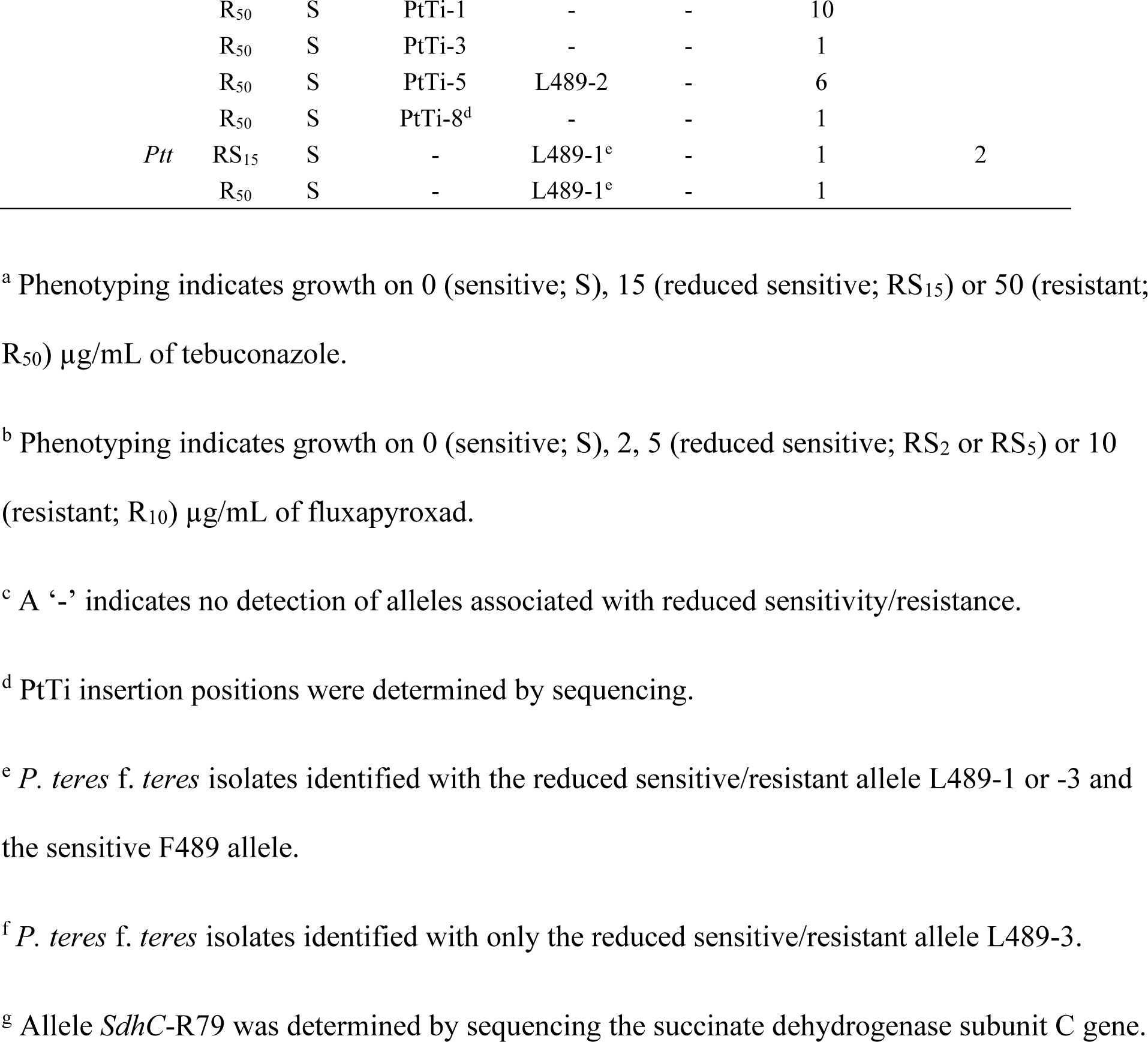
Phenotypes and genotypes corresponding to reduced sensitivity and resistance to the demethylation inhibitor (DMI) tebuconazole and succinate dehydrogenase inhibitor (SDHI) fluxapyroxad for *P. teres* f. *maculata* (*Ptm*) and *P. teres* f. *teres* (*Ptt*) isolates from the 2019 collection.

### Fungicide resistance genotypes

Real-time PCR assay development allowed genotyping for both *P. teres* f. *maculata* and *P. teres* f. *teres* and alleles associated with sensitivity, reduced sensitivity and resistance to both demethylation inhibitor and succinate dehydrogenase inhibitor fungicides (Table 1). Alignments for each assay are provided in Supplementary Figures 1 to 5.

Eight *P. teres* f. *maculata* isolates exhibited a reduced sensitivity or resistance phenotype on tebuconazole, and one *P. teres* f. *teres* isolate on fluxapyroxad, with no associated allele detected. Sequencing of the *Cyp51A* promoter region in the eight isolates determined the presence of the PtTi insertion in three positions, named as PtTi-6, PtTi-7 and PtTi-8 (GenBank Accessions OP753350, OP753351 and OP753352), and located at position −66, −69 and −89, respectively. Location positions follow the format used by Mair et al. (2020). PtTi-6 was in the same position as PtTi-4, however it contained a polymorphism which prevented the probe from binding. Geographically, PtTi-6 (*n* = 1) came from South Australia, while PtTi-7 (*n* = 5) and PtTi-8 (*n* = 2) were from Western Australia. Sequencing of *SdhB, C* and *D* sub-units (GenBank Accessions OP753347, OP753348 and OP753349) for the one *P. teres* f. *teres* isolate reported allele R79 in the *SdhC* sub-unit (C-G79R substitution; GenBank Accession OP753348). This isolate originated from South Australia.

No isolate from NSW had alleles associated with reduced sensitivity or resistance, while two *P. teres* f. *teres* from Victoria had the *Cyp51A* allele L489-3, with no detection of F489 (Table 2). For South Australia, individual *P. teres* f. *maculata* isolates were detected with *Cyp51A* alleles L489-1 (8%), L489-3 (13%) or PtTi-6 (4%). For *P. teres* f. *teres*, isolates were detected with either single or dual *Cyp51A* alleles, including L489-1 (5%), L489-3 (3% with F489 and 13% with no detection of F489), *SdhD*-Y134 (14%), *SdhC*-R134 (4%), *SdhC*-R135-1 (1%), *Cyp51A* L489-3 (with F489) + *SdhC*-R134 (46%), *Cyp51A* L489-3 (with F489) + *SdhD*-G145 (2%) or *Cyp51A* L489-3 (with F489) + *SdhC*-R79 (1%). The distribution of these alleles across two broad geographic regions (Figure 2) indicated that *SdhD*-Y134 was localised around Tumby Bay. In the region centred on the Yorke Peninsula, *Cyp51A* L489-3 (with F489) + *SdhC*-R134 was the most common genotype, with a diverse collection of genotypes present in the area.

**Figure 2.**
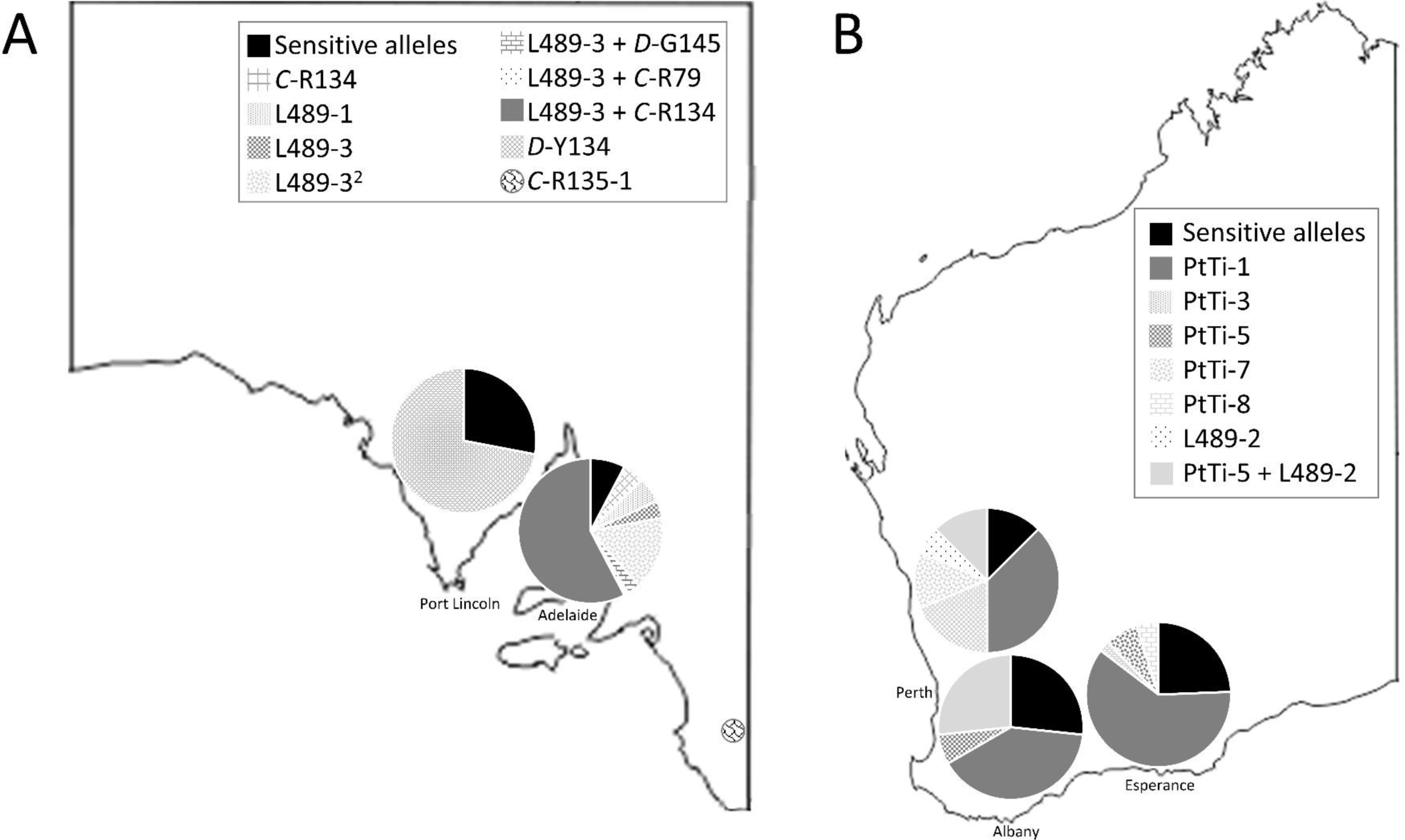
Distribution and relative frequency of fungicide sensitivity and reduced sensitivity/resistance alleles. Two geographic groups are indicated for *P. teres* f. *teres* in South Australia (A), while three broad geographic groups are indicated for *P. teres* f. *maculata* in Western Australia (B). The L489-3 allele group consists of isolates which had the L489-3 and F489 (sensitive) alleles, while the L489-3^2^ group consisted of isolates with only the L489-3 allele. The group for allele *SdhC*-R135-1 consists of a single isolate.

For Western Australia, individual *P. teres* f. *maculata* isolates were detected with *Cyp51A* allele L489-2 (1%), PtTi insertion (68%) or both alleles as L489-2 + PtTi-5 (8%). For the PtTi insertion positions, PtTi-1 was the most prevalent and was detected across three broad geographic regions (Figure 2). The L489-2 + PtTi-5 genotype was only detected in the western and northern regions, where it was the second or third most common resistant genotype, respectively. The two *P. teres* f. *teres* from Western Australia had *Cyp51A* allele L489-1.

### Comparison of fungicide resistance phenotypes and genotypes

Detection of a reduced sensitivity or resistance allele was associated with a corresponding reduced sensitivity or resistance phenotype, except for *SdhC* allele D-Y134. Of the 18 isolates with allele D-Y134, only three exhibited growth on 2 or 5 µg/mL of fluxapyroxad, with the remaining 15 reported as sensitive to fluxapyroxad (Table 2). This suggests the D-Y134 allele may confer a level of sensitivity which is below the range of the discriminatory doses. Variation in reduced sensitivity or resistance phenotypes for tebuconazole and fluxapyroxad was also present across the different alleles (Table 2).

## Discussion

This study reports a process for describing fungicide resistance phenotypes and genotypes using a combination of discriminatory fungicide doses and a comprehensive set of genotypic tests for specific alleles associated with DMI and SDHI sensitivity, reduced sensitivity or resistance. The workflow was successfully applied to net blotch samples and demonstrated the diversity of phenotypes and genotypes in barley growing regions of Australia.

Decreased sensitivity to DMI fungicides was widespread, being found in South Australia, Victoria and Western Australia. This may be due to decreased sensitivity being present in Australian populations of *P. teres* f. *maculata* and *P. teres* f. *teres* since at least 2016 and 2013, respectively (Mair et al. 2016; Mair et al. 2020). In contrast, decreased sensitivity to SDHI fungicides was first reported in 2020 for *P. teres* f. *maculata* and 2019 for *P. teres* f. *teres* in Australia (Mair et al. 2023), and was only detected in the South Australian collection in the current study. Alleles associated with DMI and SDHI reduced sensitivity or resistance varied within populations and among states, suggesting population differentiation for fungicide resistance. This is supported by Dahanayaka et al. (2021), who reported that markers aligned to the *Cyp51A* gene were significantly associated with genetic clusters in a *P. teres* f. *teres* collection. Further information on population dynamics would ideally include frequencies of reduced sensitivity or resistance alleles, however as in Mair et al. (2020), the collection procedure in the current study was not random, and true frequencies cannot be determined. Instead, the most common alleles can be suggested, which for DMI fungicides include L489-3 in *P. teres* f. *teres* from South Australia and PtTi-1 in *P. teres* f. *maculata* from Western Australia.

The detection of a single *P. teres* f. *teres* isolate with the *SdhC-*G79 allele is in contrast with reports from Europe, where G79 was the most frequently detected allele in 2013 and 2014, followed by *SdhB*-Y277 and *SdhC*-R134 alleles (Rehfus et al. 2016). In the Yorke Peninsula region of South Australia, the most common allele was *SdhC*-R134 (Mair et al. 2023), which was frequently found in dual resistance isolates with the DMI reduced sensitivity *Cyp51A* allele L489-3. Interestingly, dual resistance isolates were only *P. teres* f. *teres*, and were always detected with *Cyp51A* alleles F489 and L489-3. *P. teres* f. *teres* has two copies of the *Cyp51A* gene (Mair et al. 2016), which explains the detection of two different alleles. However, *P. teres* f. *teres* isolates with only L489-3 never had dual resistance. A distinct difference occurred in the Tumby Bay region, where the only reduced sensitivity allele was *SdhD*-Y134. The detection of six different reduced sensitivity and resistance alleles in the Yorke Peninsula region compared to one unique allele in the Tumby Bay region suggests limited spread of the pathogen between these areas. The distribution and frequency of some alleles or allele combinations may be due to a fitness cost, the impact and spread of a founder population (for example infested seed), or parallel mutation events in different fields. Each of these aspects of the pathogen’s biology require further investigation.

In a field context, changes in fungicide sensitivity may be inferred by reduced efficacy of treatments for controlling disease, or in severe circumstances, complete failure to control disease. These observations may be affected by various factors, such as non-optimal chemical application, which means laboratory confirmation of fungicide sensitivity is required (Corkley et al. 2022). *In vitro* assessments have provided the foundation for reporting reduced sensitivity or resistance emerging in *P. teres* to DMI (Campbell and Crous 2002; Ellwood et al. 2019; Mair et al. 2016; Mair et al. 2020; Sheridan et al. 1985; Sheridan and Grbavac 1985), SDHI (Mair et al. 2023; Rehfus et al. 2016) and QoI fungicides (Marzani et al. 2013; Semar et al. 2007; Sierotzki et al. 2007). The phenotyping workflow described here was designed to efficiently report the presence of reduced sensitivity or resistance to DMI and SDHI fungicides *in vitro*, however the process still required multiple culturing steps taking at least two weeks to complete. This timeframe is not conducive for in-season field management decisions. The benefits of the phenotypic test include targeted selection of *P. teres* with reduced sensitivity or resistance from leaf lesions with potentially mixed populations, reducing the chance of isolating sensitive types. Practical inclusions of multi-well plates increased the throughput of samples. A limitation of the discriminatory dose method was observed for allele *SdhD*-Y134, which was associated with sensitivity for 15 isolates. The 50% effective concentrations reported for *SdhD*-Y134 on fluxapyroxad (0.082 to 0.104) were lower than the other *Sdh* alleles associated with resistance or reduced sensitivity (0.7 to 1.75) (Mair et al. 2023). However, the 50% effective concentrations of isolates with sensitive alleles ranged from 0.006 to 0.018. This suggests the fluxapyroxad concentrations were inhibiting growth of isolates with the *SdhD*-Y134 allele in the discriminatory dose method. Instances of different phenotypes for the same genotype were also observed (for example L489-3 or *SdhC*-R134), which may have been due to variable fitness and growth rates among isolates, or procedural effects. However, with the exception of *SdhD*-Y134, the method reported a reduced sensitivity or resistance phenotype for every isolate with an associated allele. Other limitations included variable field sampling procedures and a preferential focus on reduced sensitive or resistant types. A standardised field sampling method should be considered in future studies to enable comparisons between fields and regions.

With the exception of *SdhD*-Y134, the genotyping workflow reported consistent links between the phenotypes and genotypes across the isolate collection, and resulted in the detection of four new genotypes in Australia. These were the *SdhC*-R79 allele, and three new PtTi insertion positions (PtTi-6, −7 and −8) in the *Cyp51A* promoter, including the first report of this insertion type outside of Western Australia. Future studies may be modified to initially analyse DNA from lesion tissue to detect target alleles, which has the potential to take just hours (Dodhia et al. 2021). Genetic diagnostics rely on comprehensive background knowledge of sequences associated with reduced sensitivity or resistance, and may be more technically demanding or limited when multiple sequence changes exist. For example, some assays in this study (such as *SdhC*-N75/S75) were designed for *P. teres* f. *maculata* sequences in Australia, and it was observed that these did not detect the target genes in the opposite form due to sequence differences. At this time no *P. teres* f. *teres* has been reported with the *SdhC*-S75 allele, so an equivalent assay was not designed. However, this highlights the need for continued phenotypic monitoring of populations to report the emergence of novel sequences associated with resistance.

The tools described in this study begin to address the need for information on the status of fungicide resistance in the field. PCR is likely to remain the most effective detection method for some time, however innovative approaches to collect information relevant for management decisions need to be devised. With an ability to measure frequencies of reduced sensitivity and resistance alleles, the impact of management strategies, such as fungicide rotations, could be investigated and presented to industry to allow more informed disease management.

## Acknowledgements

Isolates and leaf samples were kindly provided by Steven Simpfendorfer (Department of Primary Industries, NSW), Tara Garrard and Hugh Wallwork (South Australian Research and Development Institute, South Australia), Mark McLean (Agriculture Victoria), Andrea Hills and Geoff Thomas (Department of Primary Industries and Regional Development, Western Australia), as well as growers and agronomists. A portion of this work was performed by K. C. Adhikari as part of a Master’s project at Curtin University. This study was conducted at the Centre for Crop and Disease Management, a joint initiative of Curtin University and the Grains Research and Development Corporation (research grant CUR00023).

## Supplementary Tables and Figures

**Supplementary Table 1.**
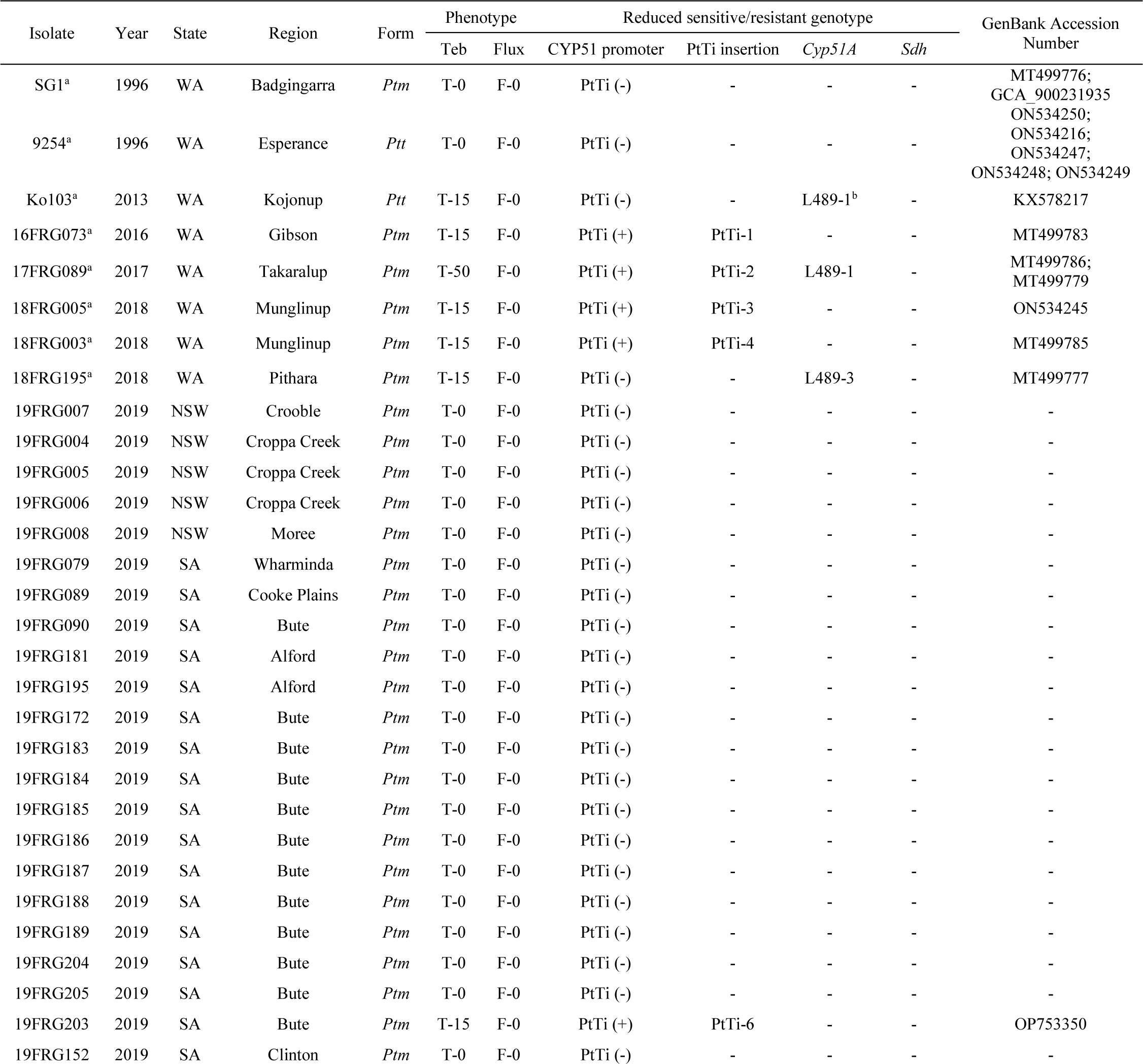
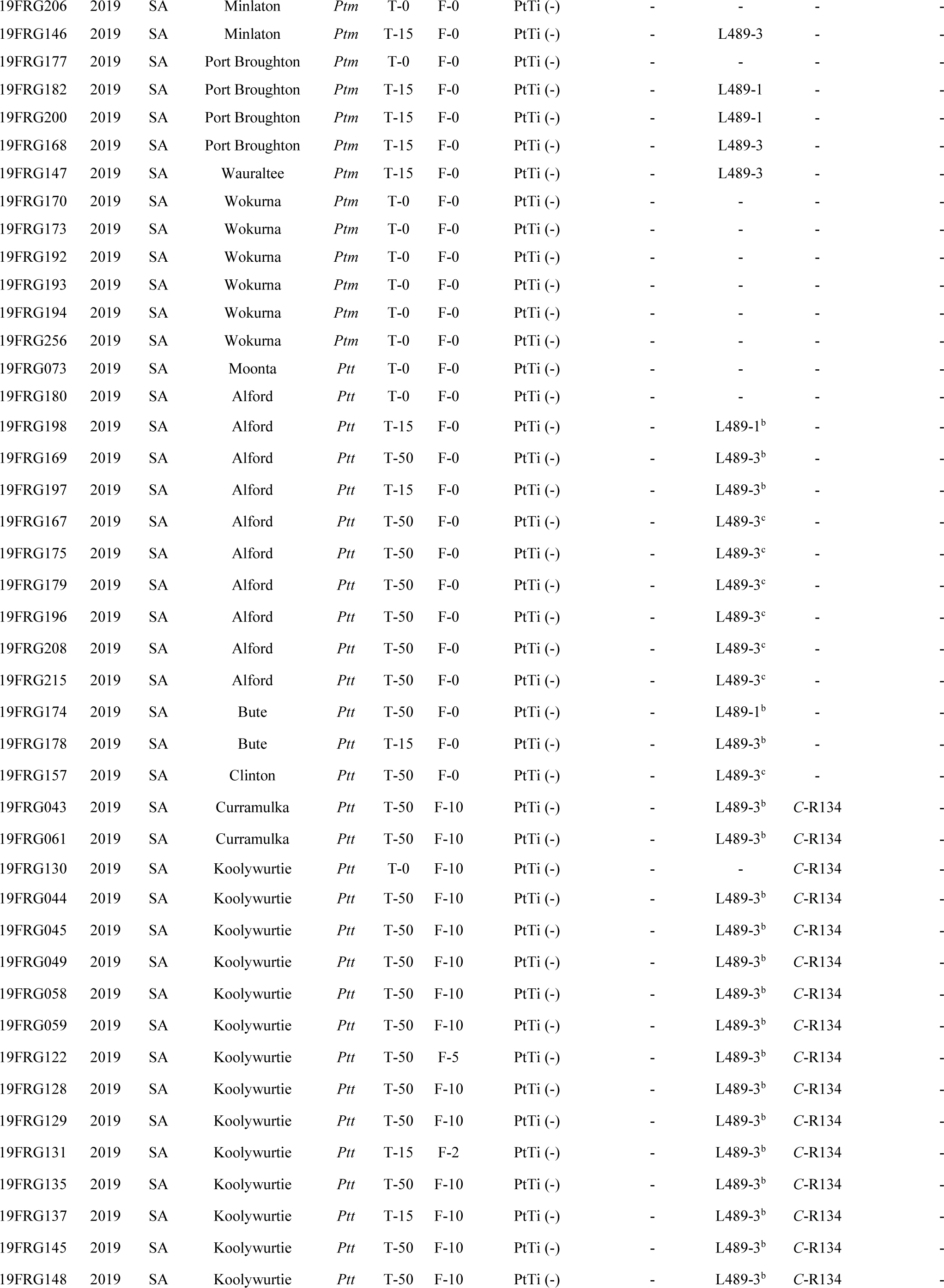
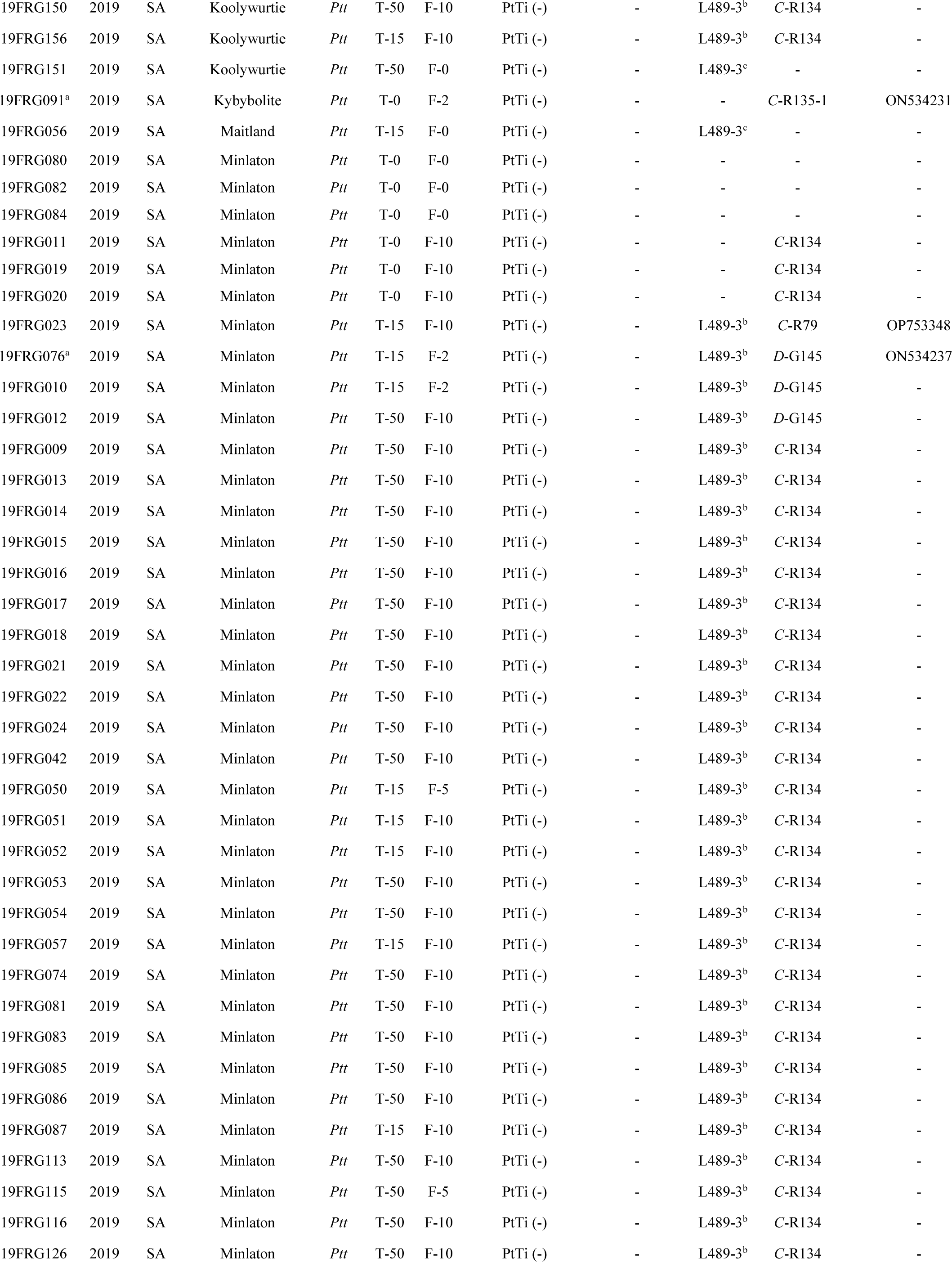
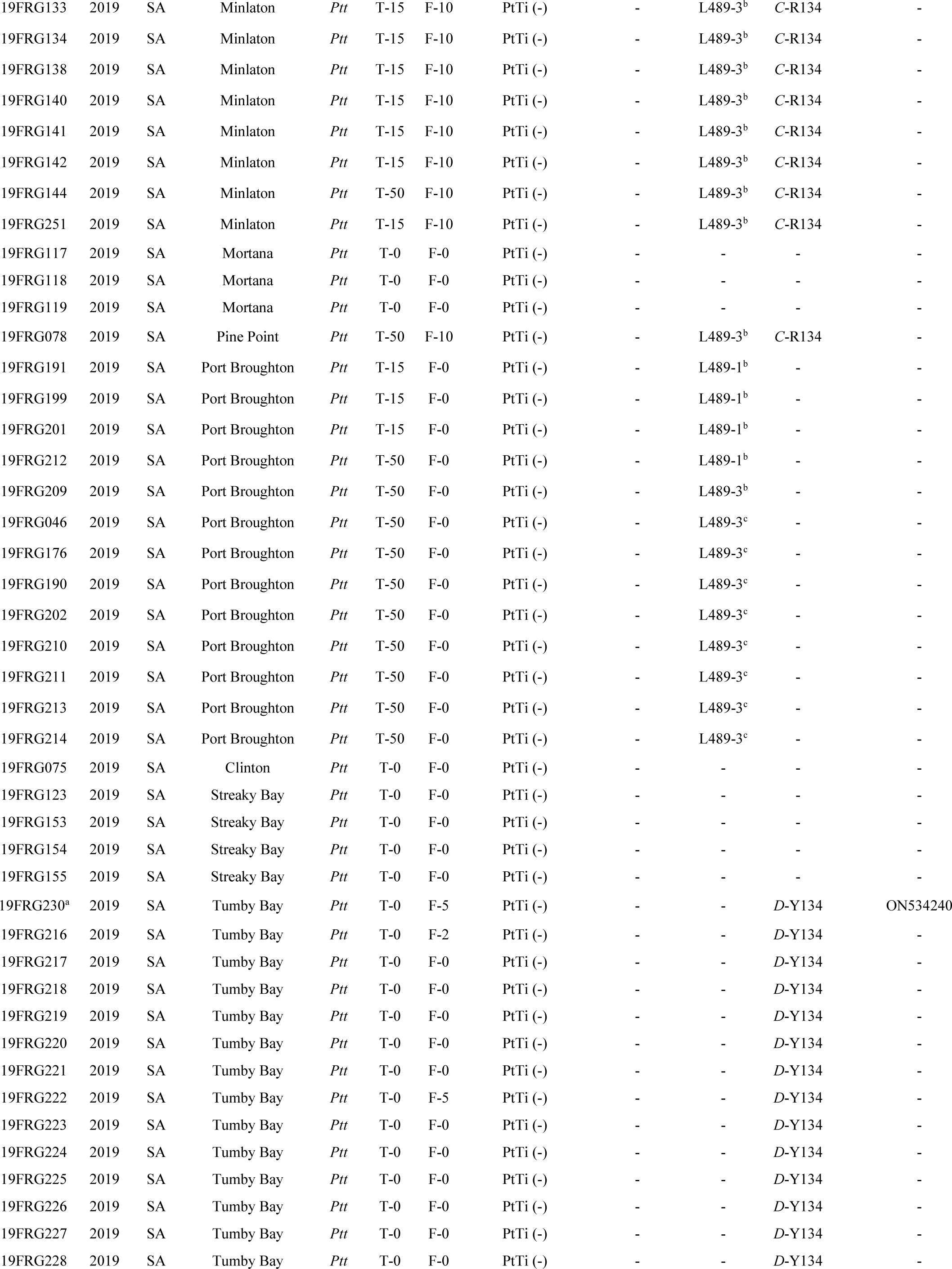
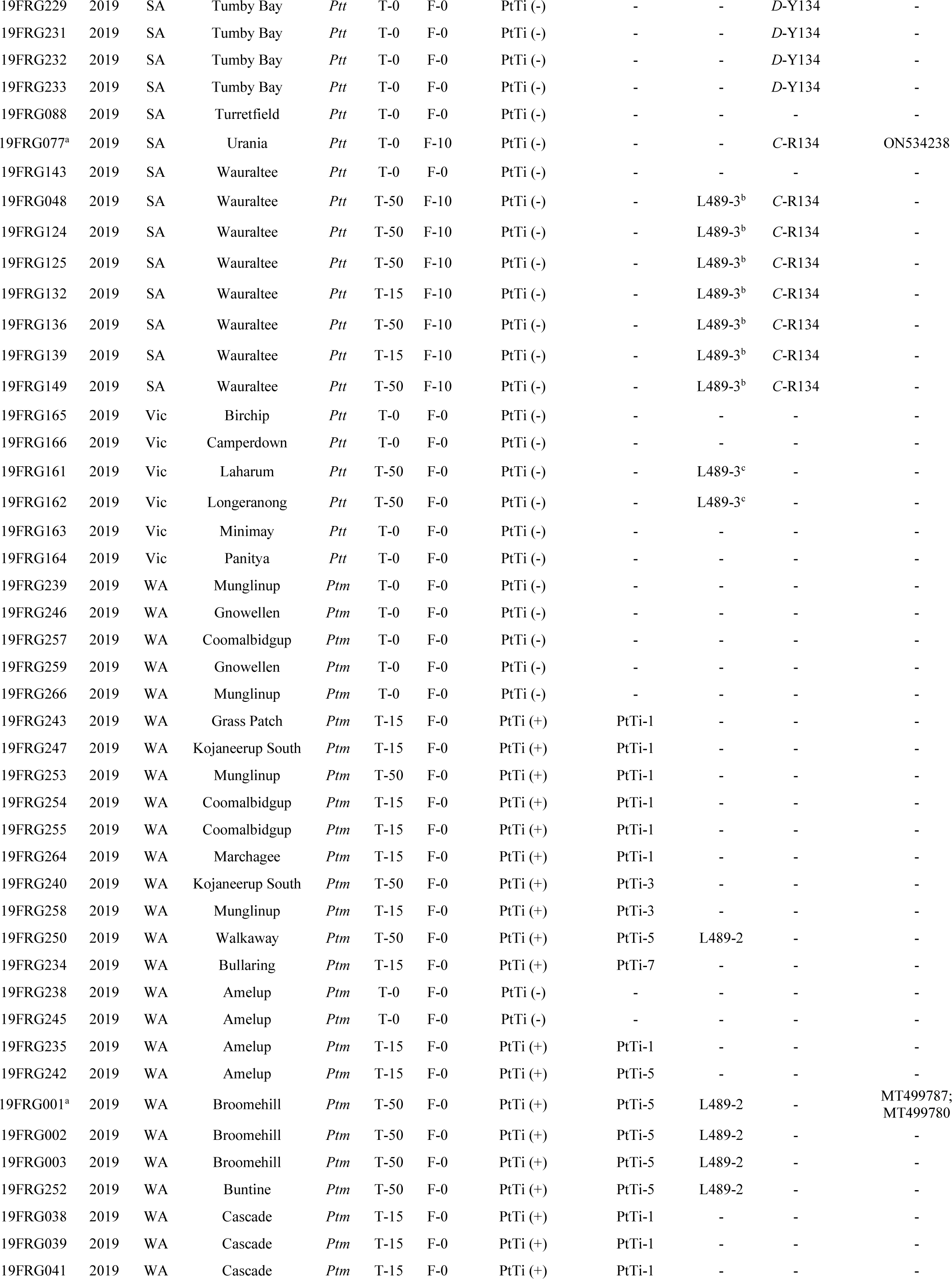
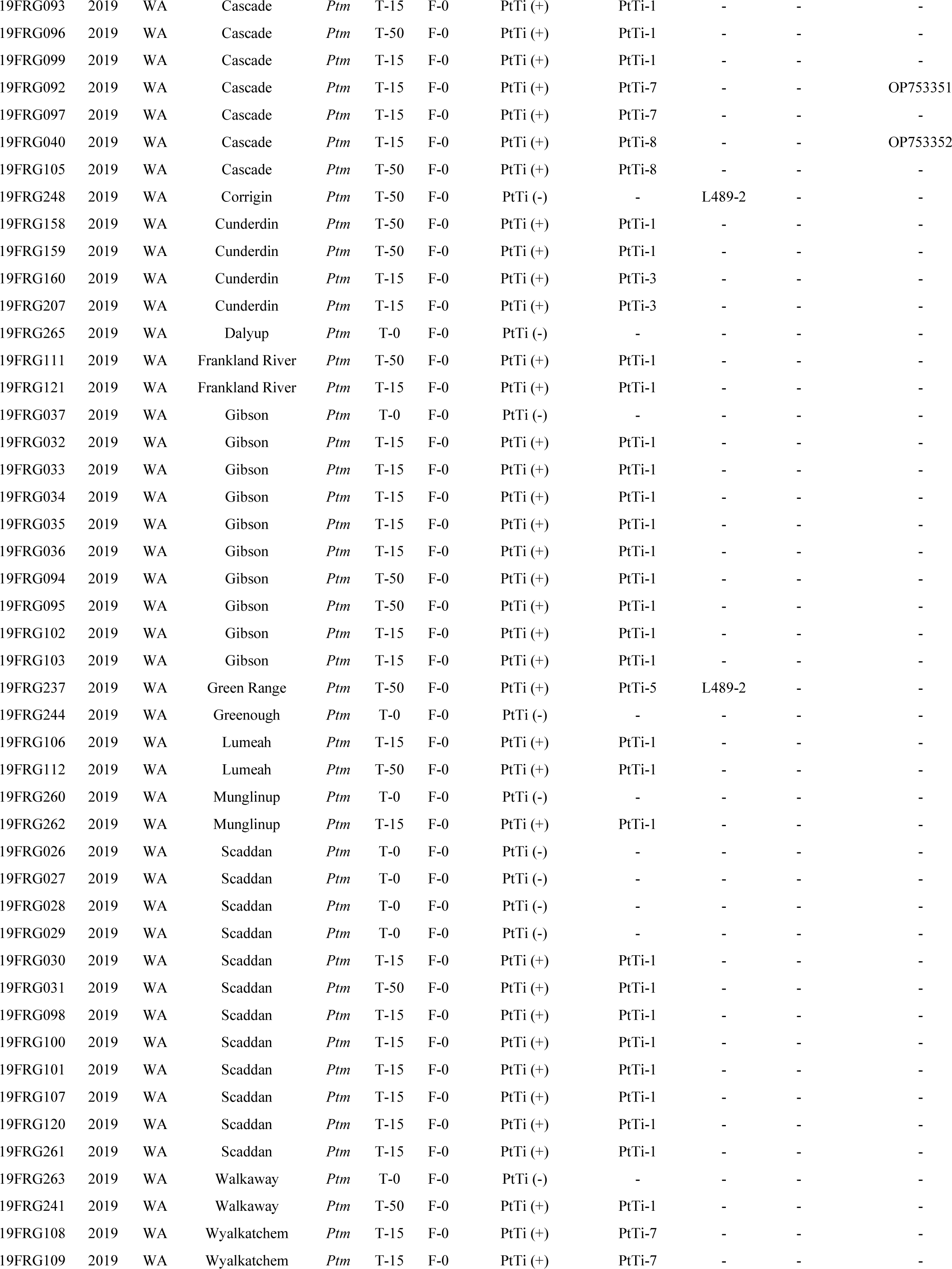
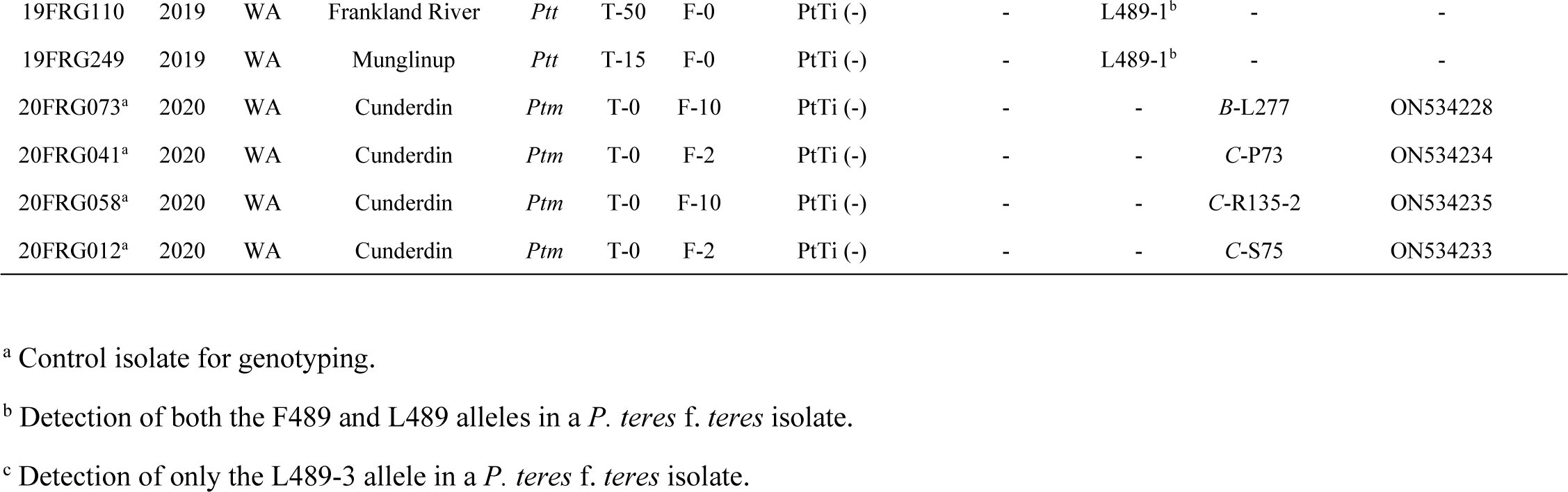
Metadata for the isolate collection assessed with phenotyping and genotyping methods. The *P. teres* f. *maculata* (*Ptm*) or *P. teres* f. *teres* (*Ptt*) form was indicated by genotyping. Phenotyping indicates growth on 0, 15 or 50 µg/mL of tebuconazole (T; demethylation inhibitor) and 0, 2, 5 or 10 µg/mL fluxapyroxad (F; succinate dehydrogenase inhibitor). Genotyping indicates the detection of alleles associated with reduced sensitivity or resistance.

